## Supplemental Information for "Light sets the brain’s daily clock by regional quickening and slowing of the molecular clockworks at dawn and dusk"

### Figure Supplements

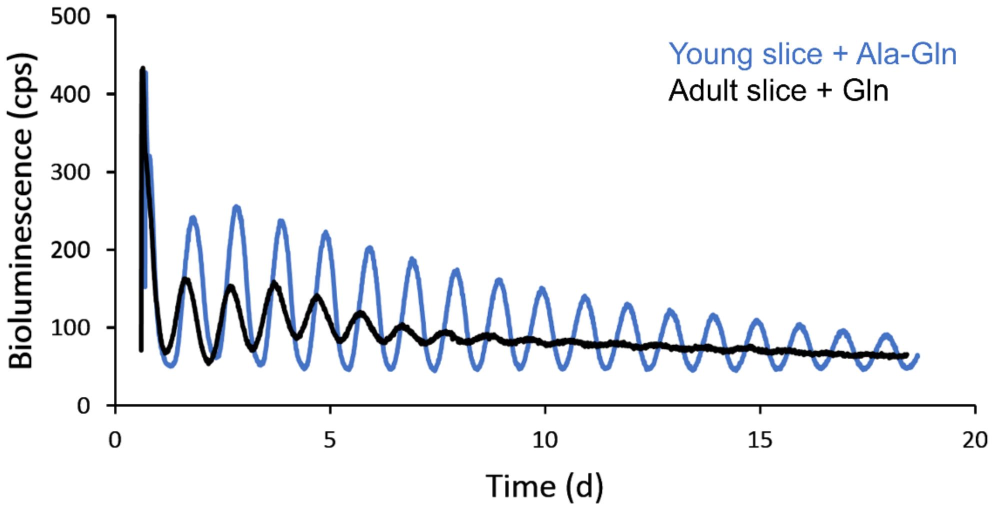


**Figure 1–figure supplement 1.** Improved PER2::LUC rhythmicity in SCN slice explanted from young mice to culture medium containing stabilized glutamine. Representative PER2::LUC rhythms of SCN slice cultures from an adult (P60) and a young (P12) mouse. Young SCN slice cultures with stabilized glutamine (alanyl-glutamine) showed higher amplitude in PER2::LUC rhythms for a longer duration, compared to rhythms from adult SCN slices in culture medium containing regular glutamine.

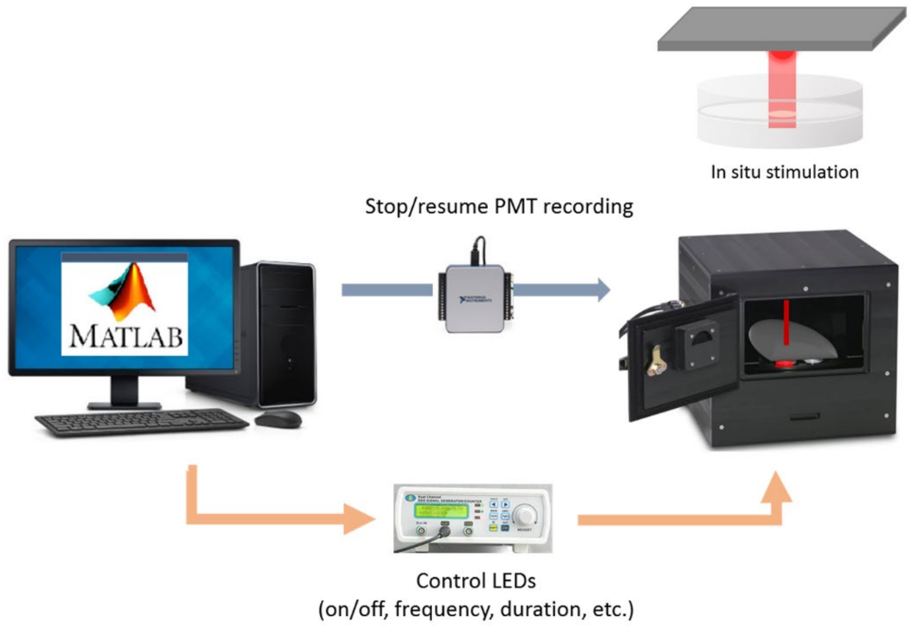


**Figure 1–figure supplement 2.** Diagram of an integrated system for long-term luminescence recording and optogenetic stimulation. Custom-written Matlab code has access to a luminometer data collection software, a multifunction I/O device turning on/off the photomultiplier tubes (PMTs), and a signal generator controlling LEDs. Thus, it can schedule periodic stimulation and execute a series of events during optogenetic stimulation — pause PMT recording, target positioning, LED stimulation, and PMT recording resumption.

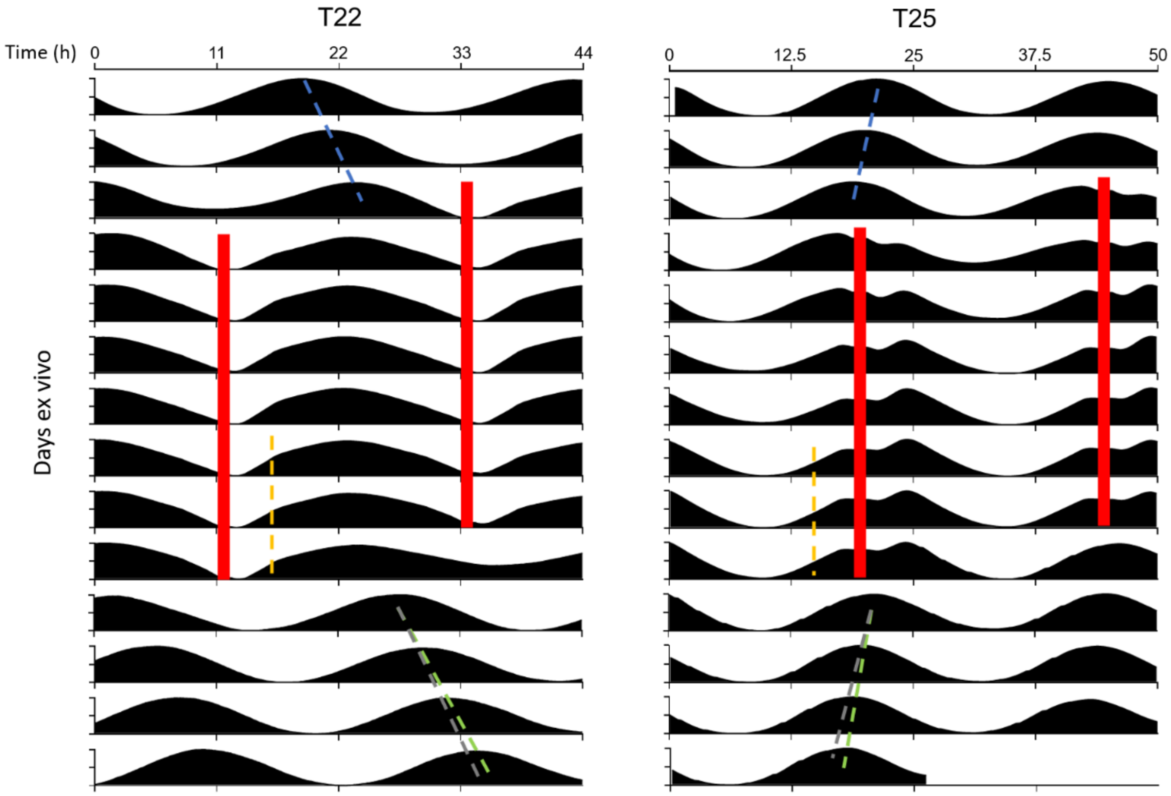


**Figure 3–figure supplement 1.** PER2::LUC rhythms in SCN slices entrain to optogenetic T-cycles. Representative double-plotted PER2::LUC bioluminescence actograms of the SCN slice entrained with 1-1.5h 10Hz optogenetic pulse (red bar) in every 22h (left) or 25h (right). Actograms are plotted against a 22h (left) or 25h (right) time scale. Linear regressions of the pre- and post-entrainment cycle peaks are indicated as the blue and green dashed lines, respectively. Yellow dashed lines indicate half-maxes on the rising phase during entrainment. Grey dashed lines indicate the pre-entrainment cycle period as a reference.

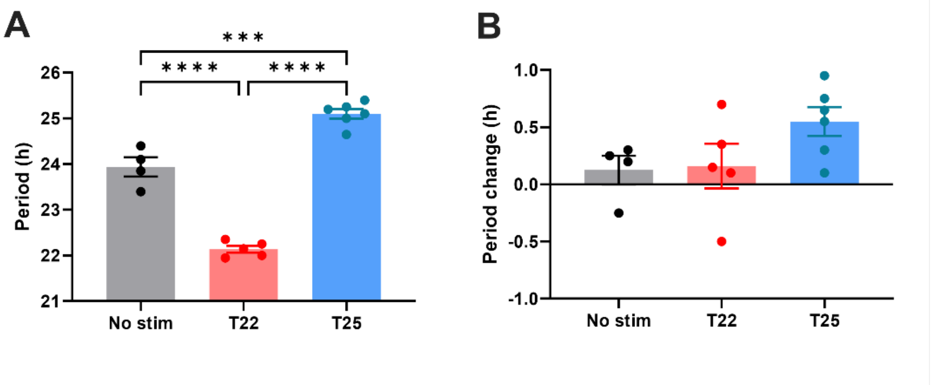


**Figure 3–figure supplement 2.** Quantification of period during entrainment (left) and period changes by entrainment (right) using Lomb-Scargle periodogram. (One-way ANOVA with Tukey's multiple comparisons tests, mean ± SEM, n = 4-6, ****$p < 0.0001$, ***$p < 0.001$)

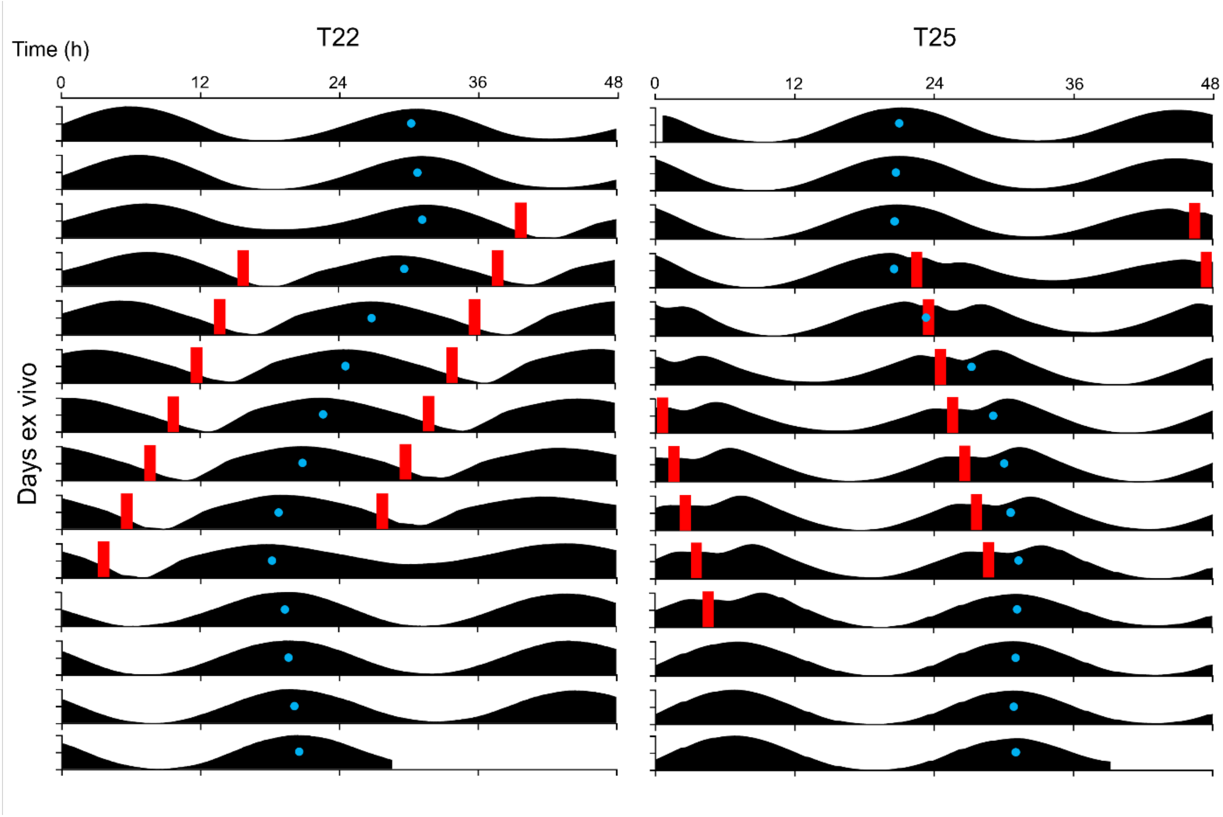


**Figure 3–figure supplement 3.** Acrophase fitting of PER2::LUC bioluminescence actograms in the Figure 3A. Blue dots denote acrophases.

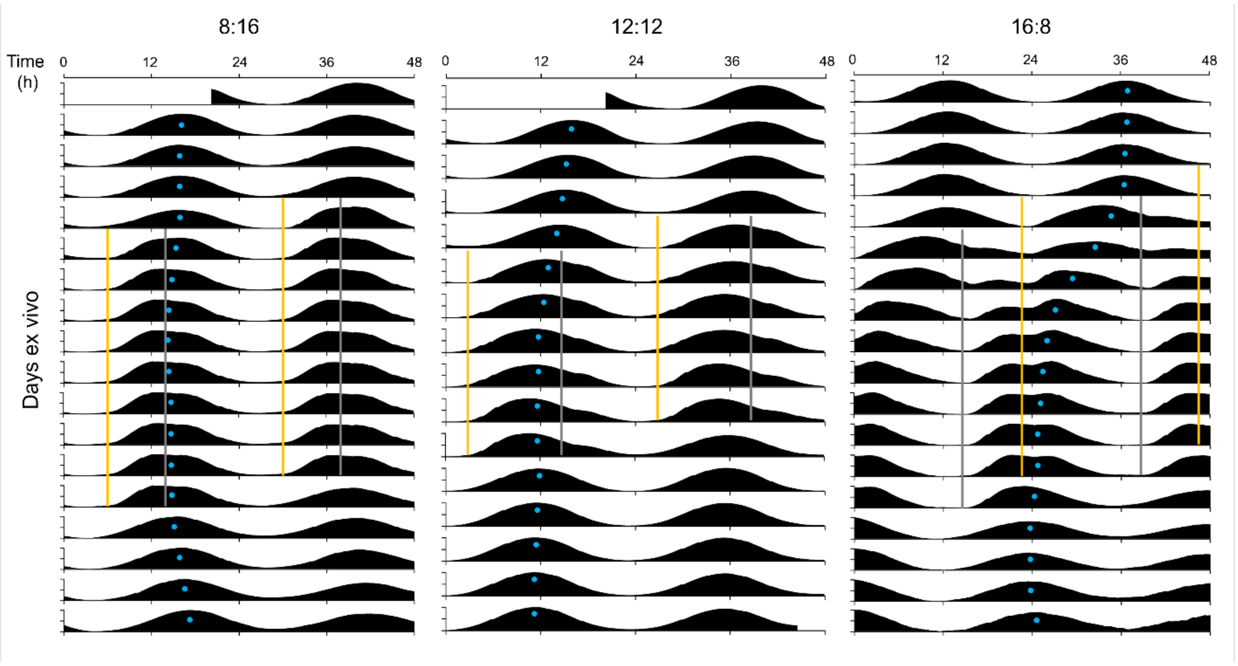


**Figure 4–figure supplement 1.** Acrophase fitting of PER2::LUC bioluminescence actograms in the Figure 4B. Blue dots denote acrophases.

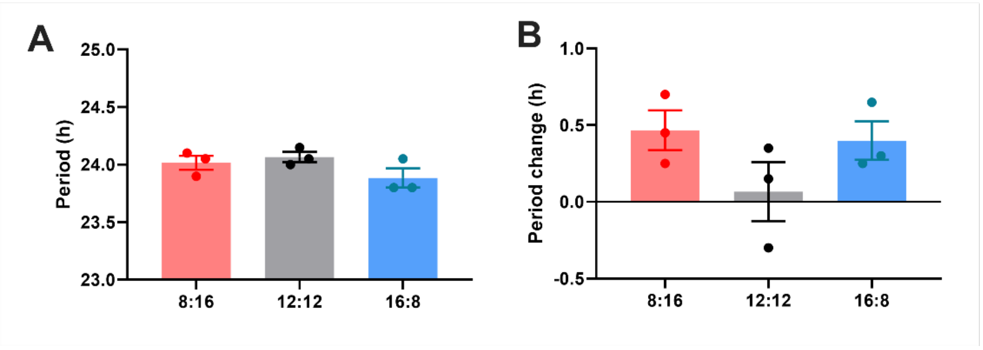


**Figure 4–figure supplement 2.** Quantification of **(A)** period during entrainment and **(B)** period changes by entrainment using Lomb-Scargle periodogram.

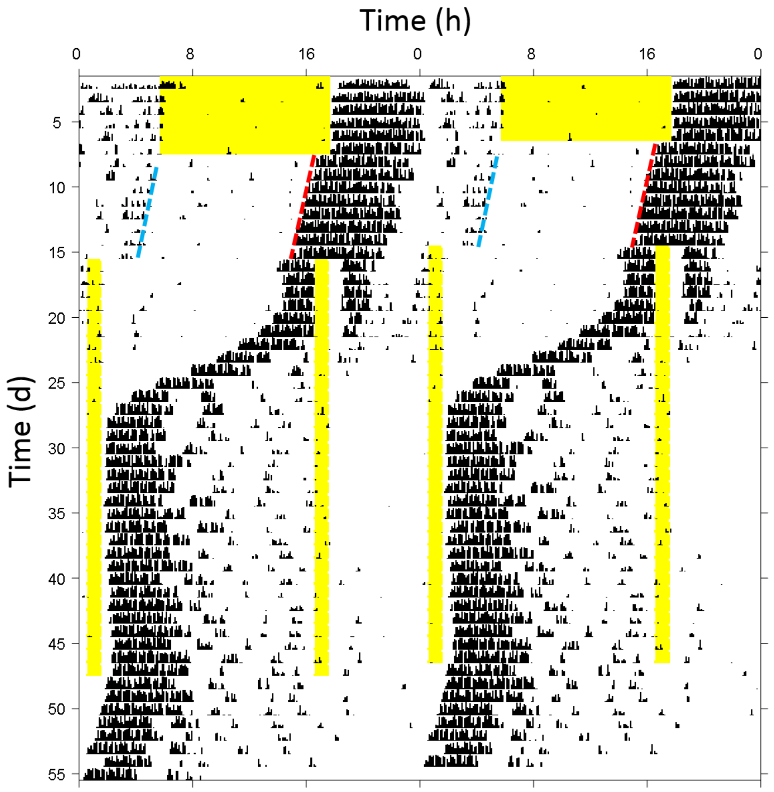


**Figure 4–figure supplement 3.** Phase jump in circadian behavior during long skeleton photoperiod entrainment. Representative double-plotted wheel-running actogram showing a behavioral phase jump to the preferred phase angle of 16:8 long skeleton photoperiod entrainment. The black tick marks indicate 6 min-binned wheel-running activity. A mouse under a light cycle of 12h light (yellow bar) and 12h darkness was released into constant darkness and then presented with a 16:8 skeleton photoperiod. The 16h interval between 1h light pulses defining dawn and dusk was initially aligned with the subjective day (i.e., the time from the nocturnal behavioral offset (blue dashed line) to the onset (red dashed line)). As the circadian rhythm rapidly phase-advanced across the nominal dusk pulse, the shorter 8h interval became aligned with the subjective day instead of the night.

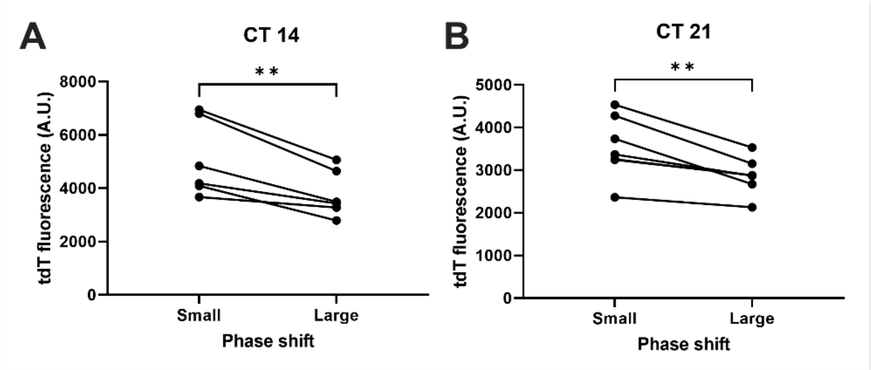


**Figure 6–figure supplement 1.** ChrimsonR-tdT fluorescence levels in different phase shift clusters in the Figure 6C for **(A)** CT14 and **(B)** CT21 stimulation. (Paired t-test, n=6-7, **p<0.01).
